# Highly pathogenic avian influenza (H5N1) landscape suitability varies by wetland habitats and the degree of interface between wild waterfowl and poultry in India

**DOI:** 10.1101/2020.09.23.310888

**Authors:** Michael G. Walsh, Siobhan M. Mor, Shah Hossain

## Abstract

Highly pathogenic avian influenza (HPAI), subtype H5N1, constitutes one of the world’s most important health and economic concerns given the catastrophic impact of epizootics on the poultry industry, the high mortality attending spillover in humans, and its potential as a source subtype for a future pandemic. Nevertheless, we still lack an adequate understanding of HPAI H5N1 epidemiology and infection ecology. The nature of the wild waterfowl-poultry interface, and the sharing of diverse wetland habitat among these birds, currently underscore important knowledge gaps. India has emerged as a global hotspot for HPAI H5N1, while also providing critical wintering habitat for many species of migratory waterfowl and year-round habitat for several resident waterfowl species. The current study sought to examine the extent to which the wild waterfowl-poultry interface, varied wetland habitat, and climate influence HPAI H5N1 epizootics in poultry in India. Using World Organisation for Animal Health reported outbreaks, this study showed that the wild waterfowl-poultry interface and lacustrine, riparian, and coastal marsh wetland systems were strongly associated with landscape suitability, and these realtionships varied by scale. Although increasing poultry density was associated with increasing risk, this was only the case in the absence of wild waterfowl habitat, and only at local scale. In landscapes increasingly shared between wild waterfowl and poultry, suitability was greater among lower density poultry, again at local scale only. These findings provide further insight into the occurrence of HPAI H5N1 in India and suggest important landscape targets for blocking the waterfowl-poultry interface to interrupt virus transmission and prevent future outbreaks.

## Introduction

Highly pathogenic avian influenza (HPAI), subtype H5N1, presents one of the biggest threats to the agricultural economy and human health. The economic burden associated with outbreaks in poultry is quick and catastrophic due to both the mortality experienced by infected birds and the standard practice of culling to prevent further spread(1–3). The threat to human health is also substantial given the realised high fatality associated with spillover, which is approximately 60% for HPAI H5N1(4). Further, through the ongoing processes of influenza A mutation and reassortment and ever-present evolutionary pressure toward novel subtypes, HPAI H5N1 poses a significant risk as a progenitor for a future pandemic subtype.

The global significance of HPAI H5N1 notwithstanding, the infection ecology of viral transmission remains poorly understood, which confounds optimal prevention strategies. Wild waterfowl, primarily Anatidae, are natural reservoirs of avian influenza A viruses, including HPAI H5 subtypes(1,5). Migratory and residential wild waterfowl maintain global cycling of these viruses along with domestic poultry(5–8). Influenza A viruses infect the intestinal tract of birds such that waterborne transmission dominates their influenza epidemiology(9). Importantly, the waterfowl-poultry interface is demarcated by shared water in the landscape (Figure 1) and, therefore, may influence both spillover (transmission from wild to domestic birds) and spillback (transmission from domestic to wild birds). Nevertheless, the nature and distribution of this interface remains unknown for many regions of the world. For example, whether certain kinds of wetlands are more important than others for the waterfowl-poultry interface, and if the interface operates differently at different densities of poultry, are both critical questions that remain unanswered. Earlier reports showed that poultry outbreaks of HPAI H5N1 over a 10-year period were strongly associated with the distribution of both surface water and epizootics in wild waterfowl(10) and poultry densities(11). However, these reports did not actually quantify the waterfowl-poultry interface or attempt to explore the specific relationship between interface and HPAI outbreaks. Moreover, few specific regions of the world have been the subject of in-depth investigations of the nature of this interface, with a notable exception being the Qinghai Lake and Poyang Lake regions in China(12).

**Figure 1.**
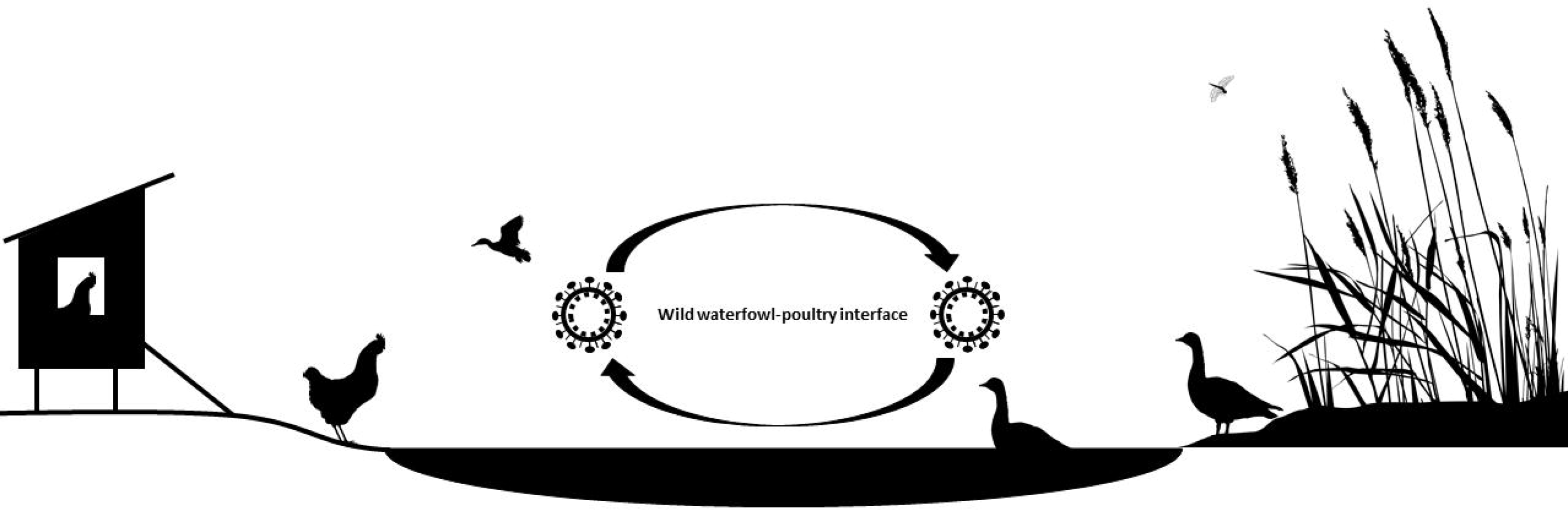
The sharing of water resources in wetland habitat is fundamental to understanding the wild waterfowl-poultry interface and its relevance to the infection ecology of highly pathogenic avian influenza (HPAI), subtype H5N1. The HPAI H5N1 virus infects, and is shed by, the intestinal tract of both waterfowl and poultry and thus the bodies of water associated with wetlands used by these birds can serve as a common vehicle for waterborne transmission in processes of both spillover and spillback.

India provides critical wintering grounds for migratory waterfowl, maintains many populations of resident waterfowl species, and is also a global hotspot for HPAI H5N1(7,8,13). Previous work has identified the importance of surface water to the landscape epidemiology of HPAI epizootics in Asia more generally(13), but did not distinguish between different wetland environments, which may facilitate waterfowl-poultry interface differently. Other work focused on Northeast India identified duck density as a key driver of HPAI H5N1 outbreaks, but did not describe the waterfowl interface or any measure of surface water(14). Similarly, a compartmental model developed by a different group suggested the effective contact rate between poultry and wild waterfowl was a key driver of outbreaks between 2008 and 2010 in the same region of India but it was limited to the single state of West Bengal(15). Nevertheless, as in many regions where HPAI presents a significant threat of spillover, the nature of the waterfowl-poultry interface has not been quantified in India nor has its influence on outbreaks been interrogated. Neither has the relative importance of migratory versus resident waterfowl to H5N1 outbreaks been explored adequately (16) despite some parts of Asia showing more than a 4-fold higher prevalence in resident waterfowl(17). The one study in India examining differences in avian influenza virus prevalence showed no difference between migratory and resident birds, but was limited by low representation of wild waterfowl in favour of larger representation of other bird families that may be less relevant as HPAI H5N1 reservoir hosts(18).

The current study sought to quantify interfaces between poultry and both resident and migratory wild waterfowl using the ecological niches of Anatidae species and subsequently evaluate their influence, and relative importance, on HPAI H5N1 outbreaks across the full extent of the diverse wetland habitats of India. Greater sharing of habitat between wild waterfowl and domestic poultry, particularly landscapes of optimal migratory waterfowl niches and low- to mid-density poultry, was hypothesised to present greater outbreak risk. This study also aimed to distinguish the influences of lacustrine (lake), riparian (river), and several types of marsh wetland environments on HPAI H5N1 outbreaks. It was hypothesised that lacustrine systems would dominate the hydrogeography of landscape suitability.

## Methods

### Data sources

Reported outbreaks of HPAI H5N1 in poultry in India between 1 January, 2006, and 30 June, 2020, and which were recorded with a precision of 5km or better, were obtained from the World Organisation for Animal Health (OIE) (n = 69). All geographic coordinates were verified in Google Maps and cross-referenced with Open Street Map to ensure that locations were correctly recorded. In some cases only coordinates for the district centroid were recorded in the reports, while specific outbreak locations were recorded as village/locality names. In these cases, the coordinates for the village/locality were obtained in Google Maps and cross-referenced with Open Street Map. While there was a clear seasonal pattern to outbreak occurrence, with high activity corresponding to the period November to April and low activity to the period May to October (Figure 2), the number of cases during the period of low activity was insufficient (n = 7) to partition the analysis into two distinct seasons. An independent set of laboratory-confirmed outbreak data derived from non-OIE sources (n = 22) was acquired through the Food and Agriculture Organisation (FAO) Global Animal Disease Information System (EMPRES-i)(19). This independent dataset was used to test the external validity of the models of HPAI H5N1 landscape suitability (see statistical methods below). These training and testing data are available at Figshare (https://figshare.com/s/10c6553ce4c73958bb9b).

**Figure 2.**
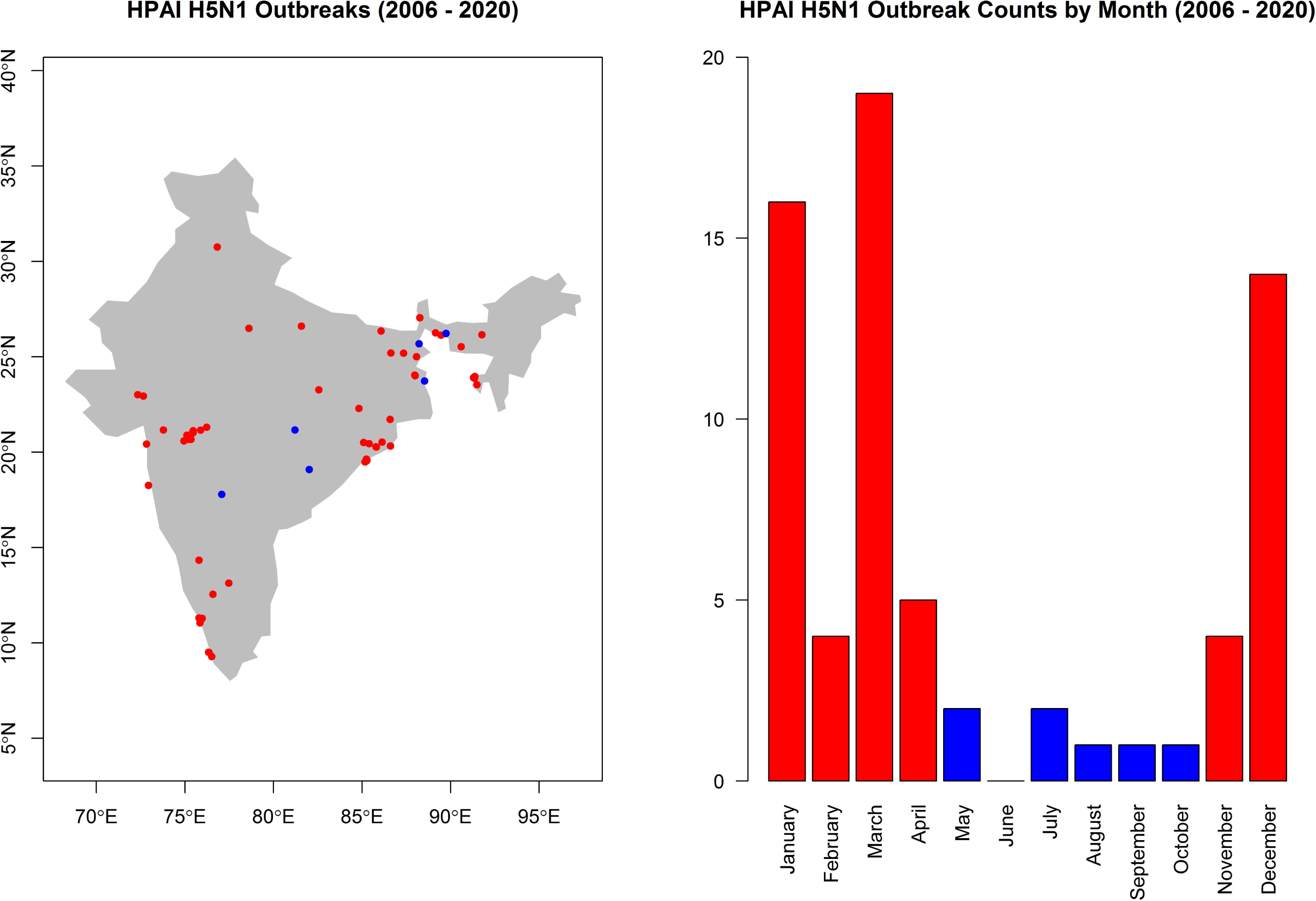
The spatial (left panel) and temporal (right panel) distributions of highly pathogenic avian influenza, subtype H5N1, outbreaks in India. Outbreaks that occurred during the high season are depicted in red and those that occurred during the low season in blue. The map is displayed only for the purposes of depicting the distribution of disease occurrence, and does not reflect the authors’ assertion of territory or borders of any sovereign country including India.

The Gridded Livestock of the World (GLW) was used to represent poultry densities for chicken and ducks(20). The 2007 edition of this data product was used to more closely correspond to the beginning of the period under which HPAI H5N1 outbreaks were observed. Although these data can manifest substantial spatial heterogeneity in error in some countries, in the Indian context poultry estimates were adjusted by animal censuses at the 2nd and 3rd stage administration levels, corresponding to the district and taluk, respectively, thereby yielding an acceptable level of data verification at a sub-state scale(20). The Global Biodiversity Information Facility (GBIF) was used to acquire all observations of Anatidae species (282,850 individual observations of 23 distinct species) between 1 January, 2000 and 31 December, 2019 across Pakistan, India, Nepal, Bhutan, and Bangladesh so the ecological niches of each species could be modelled across the region (http://www.gbif.org/). Observations in all South Asian countries were included to correct for edge effects and to better represent the environmental niches of the species involved, many of which winter or reside permanently in other parts of South Asia, rather than India alone.

To correct for potential spatial reporting bias in the observations of wild waterfowl species used in species distribution models, background points were weighted by the human footprint (HFP) (see modelling description below). The data used to represent HFP were obtained from the Socioeconomic Data and Applications Center (SEDAC) repository(21), and quantified in two steps(22). First, an index of human influence impact was constructed from eight domains: 1) population density, 2) proximity to roads, 3) proximity to rail lines, 4) proximity to navigable rivers, 5) proximity to coastlines, 6) spectrum of night-time artificial light, 7) urban vs rural location, and 8) land cover. The eight domains are scored and subsequently combined to create the human influence index (HII), which ranges from 0 (no human impact) to 64 (maximal impact). The final HFP metric is the ratio of the range of minimum and maximum HII in the local terrestrial biome (one of 15 terrestrial biomes defined by the World Wildlife Fund) to the range of minimum and maximum HII across all biomes, and is expressed as a percentage(22).

Surface water data were obtained from the Global Lakes and Wetlands Database(23). Hydrogeographical classifications were as follows: coastal wetland, river, controlled water reservoir, lake, freshwater marsh, swamp, or intermittent wetland(24). New distance rasters were created from each surface water type using the proximity function in the QGIS geographic information system(25). Pixel values in these rasters convey the distance in kilometres between each surface water type and all other pixels within the geographic extent under study. Thus, relationships between outbreaks and distinct surface water environments can be investigated.

WorldClim Global Climate database provided the requisite climate data for this study(26). Because of the distinct seasonal pattern in HPAI H5N1 outbreaks, the current analysis focused on seasonal measures of both temperature and precipitation. Thus, rasters for the mean warmest quarter temperature and coldest quarter temperature, and the mean driest quarter precipitation and wettest quarter precipitation, were extracted.

### Statistical Analysis

#### Wild waterfowl species distribution modelling

The environmental niches of each Anatidae species with at least 100 field observations were modelled using boosted regression trees (BRT). Under this machine learning approach, algorithms partition a data space according to rules that optimize homogeneity among predictors and a response (i.e., the presence of the species of interest). The derived data space forms decision trees(27), which are 1) robust to outliers, 2) comprised of predictors that need not adhere to the model form assumptions of generalised linear models, and 3) capable of modelling interactions between the predictors as a consequence of their hierarchical structure. The sampling of background points used in the BRT models was weighted by the human footprint to correct for potential sampling bias in field observations of waterfowl species, as spaces with greater accessibility would be expected to be more represented in the field sample for each species. Approximately 10000 regression trees maximised the fit of the BRT model for each species, while shrinkage was set to 0.001 to maximise the learning rate for a large number of trees(27,28). Models were trained using 5-fold cross-validation, with fit and performance assessed by the cross-validation error and the area under the curve, respectively. The environmental features used to model species’ distributions comprised mean precipitation during the driest and wettest quarters, mean annual temperature, isothermality, and distance to the nearest surface water. These variables exhibited low correlation with each other (all Pearson’s r < 0.5) and therefore their inclusion together in the model was justified. All species distributions were modelled at a spatial resolution of 30 arc seconds (∼1 km). Distinct niches were modelled for each of 17 different migratory species and six resident species. Individual species’ names, their number of field observations, and their niche model metrics are presented in S1 Table 1.

**Table 1.** Adjusted relative risks and 95% confidence intervals for the associations between highly pathogenic avian influenza subtype H5N1 outbreaks and each landscape feature as derived from the best fitting inhomogeneous Poisson models for resident and migratory waterfowl, respectively. Each landscape feature is adjusted for all others in each of the two models.

| Landscape feature | Relative risk | 95% confidence interval | p-value |
| --- | --- | --- | --- |
| *Model 1: with resident waterfowl* |  |  |  |
| Poultry density (deciles) | 1.29 | 1.06 – 1.57 | 0.01 |
| Resident wild waterfowl composite niche (%) | 147.43 | 7.81 – 278.40 | 0.0008 |
| Resident waterfowl:poultry interaction | 0.60 | 0.40 – 0.89 | 0.01 |
| Distance to coastal marsh (5 km) | 0.75 | 0.64 – 0.89 | 0.0007 |
| Distance to lake (5 km) | 0.03 | 0.003 – 0.24 | 0.001 |
| Distance to river (5 km) | 0.47 | 0.30 – 0.74 | 0.001 |
| *Model 2: with migratory waterfowl* |  |  |  |
| Poultry density (deciles) | 1.27 | 1.03 – 1.56 | 0.03 |
| Migratory wild waterfowl composite niche (%) | 65.86 | 4.22 – 102.8 | 0.003 |
| Migratory waterfowl:poultry interaction | 0.67 | 0.46 – 0.99 | 0.04 |
| Distance to coastal marsh (5 km) | 0.74 | 0.64 – 0.87 | 0.0002 |
| Distance to lake (5 km) | 0.03 | 0.003 – 0.26 | 0.002 |
| Distance to river (5 km) | 0.48 | 0.31 – 0.76 | 0.002 |

Subsequent to the modelling of individual species’ distributions, two composite niches were modelled for all migratory species combined and all resident species combined using a similar approach to that employed in South America(29). The summaries of these two composite niches are also presented in S1 Table 1. The degree of niche overlap(30) between each individual species distribution and their respective composite species distribution (migratory or resident) was evaluated to determine the nature and extent of the level of heterogeneity between the specifies-specific environmental niches for each of the two waterfowl groups. The objective of assessing niche overlap in this way was to determine 1) if heterogeneity was so extensive that no composite niche for migratory or resident species could be justified, 2) if a small number of species demonstrating divergent niches should be considered individually in concert with composite migratory and resident niches for the remaining species, or 3) if the species demonstrated such homogeneity in their environmental niches that only composite niches for migratory and resident species would be required to represent the distribution of wild waterfowl across India.

The gbm function in the gbm package was used to fit the BRT models(28,31) in R v. 3.6.1(32), while the dismo package was used to compare niche overlap(33).

#### HPAI H5N1 Outbreak modelling

The HPAI H5N1 outbreaks were fitted as a point process(35). Under this model framework the spatial dependence of the outbreak distributions can be determined and, where dependencies are in evidence, can be assessed with respect to environmental features.

First, HPAI H5N1 outbreaks were fitted as a homogeneous Poisson process, with conditional intensity,

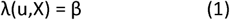

where u designates the geographic locations of outbreaks, X, and β is the intensity parameter. The intensity is defined as the number of points in a subregion of a window of a specified geographic extent. The homogeneous Poisson model represents the null model for complete spatial randomness (CSR). Under CSR, the expected intensity is simply proportional to the area of the subregion under consideration(35), i.e. does not demonstrate spatial dependency.

Second, the model under CSR was compared to an inhomogeneous Poisson process, which incorporates spatial dependency in HPAI H5N1 outbreaks and has conditional intensity,

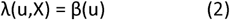

In contrast to (1), here the intensity is represented as a function of the location, u, of the HPAI H5N1 outbreaks. Outbreak intensity demonstrated spatial dependence based on the better fitting inhomogeneous Poisson process model and the significant divergence of the K-function from CSR (see results below). Bivariate and multiple inhomogeneous point process models with environmental features were fitted with conditional intensity,

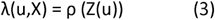

where ρ is the parameter representing the association between the point intensity and the feature Z at location u. The background points used in the model were sampled proportional to HFP, to control for potential reporting bias in the HPAI H5N1 surveillance mechanisms.

Because a primary aim of this study was to examine how HPAI H5N1 landscape suitability varies by the degree of interface between wild waterfowl and poultry, poultry density was not included as an offset but rather evaluated explicitly as a covariate in these models. Covariates were aggregated to scales of 2.5 and 10 arc minutes, respectively, for the two spatial scales under evaluation (see below). The crude associations between HPAI H5N1 outbreaks and mean dry quarter precipitation, mean wet quarter precipitation, mean warm quarter temperature, mean cold quarter precipitation, proximity to each surface water class, the composite ecological niches of resident and migratory Anatidae species, the niches of *Mergus merganser, Mareca falcata*, and *Asarcornis scutulata*, which diverged from their respective composite niches (see Results), and deciles of poultry density were first evaluated in bivariate models with a separate simple inhomogeneous Poisson model for each feature (S2 Table 2). Those features with confidence intervals that did not include 0 were included as covariates in the multiple inhomogeneous Poisson models (S3 Figure 1, S4 Figure 2, S5 Figure 3). Covariates included in the multiple inhomogeneous Poisson models demonstrated low correlation (all values of the Pearson’s r were < 0.5). The regression coefficients from the inhomogeneous Poisson models were used to compute relative risks, which represent the associations between outbreak intensity and each covariate. The wild waterfowl-poultry interfaces were evaluated separately using resident- and migratory-specific models with interaction terms included in the model structures to assess the effect modification of poultry distribution by the distributions of resident and migratory waterfowl niches, respectively. Model fit was assessed using the Akaike information criterion (AIC). Model performance was tested against an independent, laboratory-confirmed set of outbreak data derived from EMPRES-i, as described above. The use of independent data for testing model performance enhances the value of this work considerably by providing a test of the external validity of the results. Performance was evaluated using the area under the receiver operating characteristic curve (AUC). Model selection proceeded by comparing the full model to reduced model groups nested on three specific domains (climate, hydrogeography, and bird hosts), and finally to the best-fitting (lowest AIC) reduced model, which comprised elements of the different domains.

**Figure 3.**
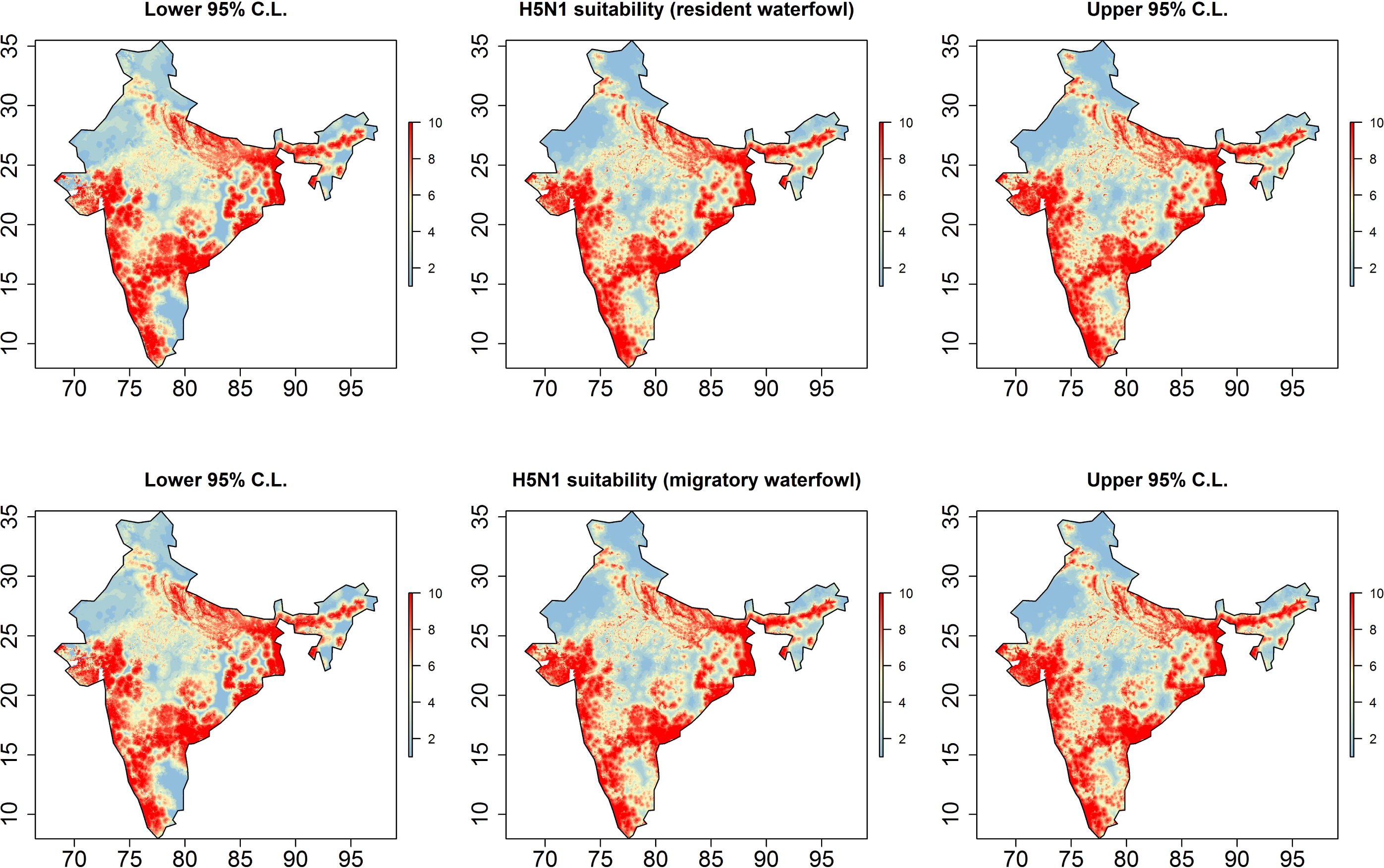
Highly pathogenic avian influenza (HPAI), subtype H5N1, landscape suitability based on predicted intensity at 2.5 arc-minutes (approximately 5 km). The centre panels depict HPAI H5N1 suitability for resident-specific (top) and migratory-specific (bottom) waterfowl models based on the predicted intensities from the best fitting and performing inhomogeneous Poisson point process models (Table 1; S6 Table 3). The left and right panels depict the lower and upper 95% confidence limits, respectively, for the predicted intensities. All maps are displayed only for the purposes of depicting the distribution of disease occurrence and risk, and do not reflect the authors’ assertion of territory or borders of any sovereign country including India.

Models were fitted and assessed at relative fine (2.5 arc-minutes) and coarse (10 arc-minutes) scale to determine if the relationships were scale-dependent since biotic (wild waterfowl and poultry) and abiotic features (hydrogeography and climate) have been shown to scale differently in some infectious disease processes(36). Again, model fit (AIC) and performance (AUC) were compared across the two scales. Spatial dependencies were identified by fitting K-functions to the HPAI H5N1 outbreaks before and after point process modelling with the specified environmental features to determine if these features adequately accounted for the observed dependencies. The R statistical software version 3.6.1 was used to perform the analyses described(32). The spatstat package was used to fit the point process models and estimate the K-functions(37,38). The silhouette images of poultry and waterfowl in Figure 1 were obtained from phylopic.org and used under Creative Commons license.

## Results

The majority of migratory waterfowl species and resident waterfowl species exhibited substantial homogeneity (niche overlap >= 90%; S1 Table 1) in their modelled environmental niches, suggesting a composite niche for each group would be justified. However, 2 migratory species, *Mergus merganser* (niche overlap = 67.8%) and *Mareca falcata* (niche overlap = 75.8%), and 1 resident species, *Asarcornis scutulata* (niche overlap = 52.1%), exhibited substantial divergence from composite migratory and resident species niches, respectively, and so were considered independently in modelling HPAI H5N1 outbreaks.

The best fitting and performing models of HPAI H5N1 landscape suitability for the resident and migratory waterfowl (Models 8 and 9, respectively, S6 Table 3) are presented in Table 1. Outbreaks were strongly associated with the niche of wild resident waterfowl species (RR = 147.43; 95% C.I. 7.81 – 278.40), poultry density (RR = 1.29; 95% C.I. 1.06 – 1.57), and their interface as represented in the model by their interaction term (RR = 0.60; 95% C.I. 0.40 – 0.89). Importantly, this negative effect modification showed that, HPAI H5N1 suitability was highest in landscapes of low-density poultry and high waterfowl suitability, while risk was also high in high-density poultry landscapes absent of wild waterfowl. Proximity to coastal marsh (RR = 0.75; 95% C.I. 0.64 – 0.89), lake (RR = 0.03; 95% C.I. 0.003 – 0.24), and river (RR = 0.47; 95% C.I. 0.30 – 0.74) wetlands were also strongly associated with suitability. While mean precipitation during the wettest quarter and mean temperature during the coldest quarter were both associated with HPAI H5N1 outbreaks in bivariate models (S2 Table 2), climate was no longer associated with suitability once the distribution of wild waterfowl, poultry, and hydrogeography were accounted for. The model with migratory waterfowl was very similar to the model with resident waterfowl, though the latter was a slightly better fit to the data at fine scale (S6 Table 3). Landscape suitability estimates for HPAI H5N1 outbreaks and their 95% confidence limits for both the resident and migratory waterfowl models are presented in Figure 3. Again, the similarity of the distribution of landscape suitability between migratory and resident waterfowl at local scale is clear. The spatial dependency identified for HPAI H5N1 outbreaks with the homogenous K-function (left panels, Figure 4) was explained by the fitted inhomogeneous Poisson models including either resident or migratory waterfowl as is shown in the right panels of Figure 4. Finally, the outbreak models assessed at the coarser scale of 10 arc minutes, showed a similar distribution of landscape suitability overall (S7 Figure 4, S8 Figure 5) and continued to show strong associations between outbreaks and both waterfowl and wetland habitats. However, there were also important differences across scale (S9 Table 4). First, the interface between waterfowl and poultry was no longer influential at this scale as suggested by the lack of association with their interaction terms in both resident and migratory waterfowl models. Second, there was no longer any difference in suitability by poultry density at this scale. Third, although the associations with both resident and migratory waterfowl remained strong, both were also considerably attenuated. Fourth, the same hydrogeographic configuration was apparent for the resident waterfowl model at this scale, whereas proximity to lakes was no longer associated with outbreaks at this scale for the migratory waterfowl model. Fifth, despite a generally similar distribution of landscape suitability for HPAI H5N1 across India assessed at 2.5 arc minutes and 10 arc minutes, there was more divergence in suitability predicted by the resident and migratory waterfowl models at 10 arc minutes (S7 Figure 4).

**Figure 4.**
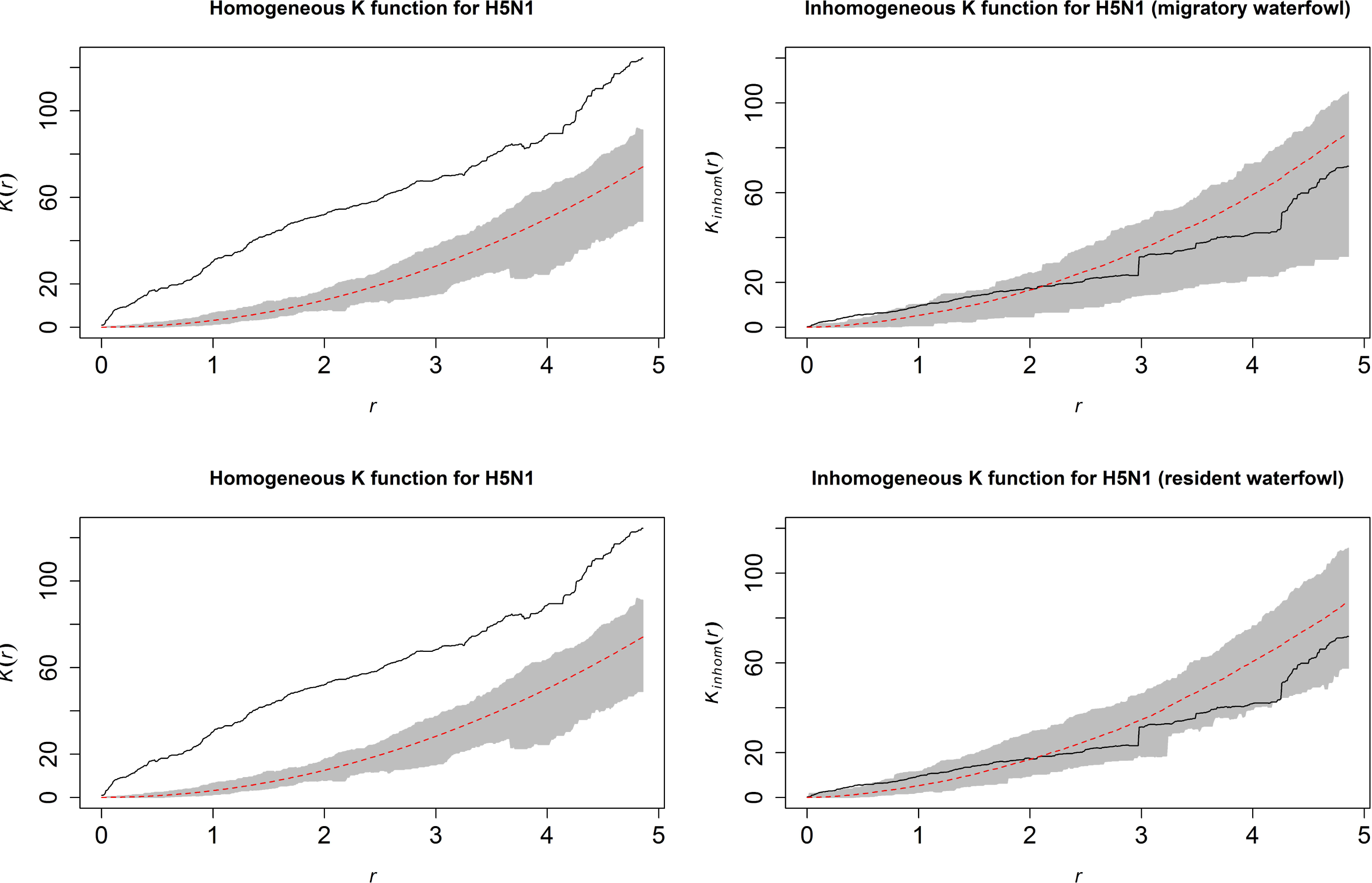
Homogeneous (left panels) and inhomogeneous (right panels) K-functions for the highly pathogenic avian influenza (HPAI), subtype H5N1, outbreak point process. The homogeneous K-function is not an appropriate fit due to the spatial dependency in HPAI H5N1 outbreaks as depicted by the divergent empirical (black line) and theoretical (under spatial randomness; dashed red line with confidence bands in grey) functions. In contrast, the resident (top) and migratory (bottom) waterfowl model-based inhomogeneous K-functions show that the spatial dependency was accounted for by the model covariates (overlapping empirical and theoretical functions). The x-axes, *r*, represent increasing radii of subregions of the window of HPAI H5N1 outbreaks, while the y-axes represents the K-functions.

## Discussion

This is the first study to systematically investigate the extent of the interface between wild waterfowl and poultry and its influence on the risk of HPAI H5N1 outbreaks in India. The current findings showed that at local scale, suitability was strongly associated with both resident and migratory wild waterfowl, their interface with poultry, and close proximity to coastal marsh, lake, and river environments. Importantly, the highest suitability corresponded to wild waterfowl interfaces with relatively low-density poultry holdings. Furthermore, the association with the wild waterfowl-poultry interface varied substantively by scale, with the influence of this interface diminishing markedly at broader scale, whereas the associations with hydrogeography were more consistent across scale, although lacustrine systems also diverged. These findings did not show a definitive distinction between the relative influences of resident and migratory wild waterfowl niches, but instead highlight more generally the importance of targeting key points of interface between wild waterfowl and domestic poultry and their wetland connections to block ongoing cycles of transmission of HPAI H5N1.

The importance of viral sharing between wild waterfowl and domestic poultry is well recognised(5–7). Moreover, genomic homogeneity of avian influenza A virus within established migratory flyways in Asia suggests the importance of migratory reservoirs to the inter-regional and inter-continental maintenance and local circulation(8,39,40). However, specific investigation of the interface between wild waterfowl and poultry in India has been limited. One study explored the impact of potential contact between wild waterfowl and poultry using discrete compartmental models, which parameterised model contact rates by empirical observations(15). While this work yielded useful results and provided some of the only insight into modelling the effect of contact between wild waterfowl and poultry on HPAI H5N1 outbreaks, its scope was limited to one part of a single state in Northeast India and it did not quantify the wild waterfowl-poultry interface more generally. The current study provides much needed context to the circulation of HPAI H5N1 in India by quantifying the interface across the region based on the ecological niche of 23 Anatidae species and their shared space with domestic poultry. Consequently, we found that the distribution of wild waterfowl and their interface with poultry, particularly at low-density poultry holdings is one of the key local drivers of outbreaks. However, landscapes with minimal wild waterfowl showed that HPAI H5N1 suitability increased with increasing poultry density, which corroborates previous studies examining risk variation across poultry density(11,14). This suggests that within the Indian context small poultry holdings drive risk in landscapes most suitable to wild waterfowl, while larger, more commercial poultry holdings drive risk in landscapes less suitable to wild waterfowl. The finding was as expected since poultry in lower density settings may experience more frequent contact with wild waterfowl via the sharing of common water resources, and also corroborates predictions from previous localised deterministic modelling(15). Interestingly, the association with the waterfowl-poultry interface was much attenuated at broader scale, despite wild resident and migratory waterfowl niches remaining associated with outbreaks. This would seem to suggest that the waterfowl-poultry interface (i.e. potential inter-specific interaction) dominates outbreak risk in local, but not regional, contexts, even though wild waterfowl niches continue to exert an attenuated influence at broad scale. The latter is not unexpected given that the influence of the niche would be expected to operate to some degree by way of the common vehicle of water, which also maintained strong influence at broad scale as coastal marsh and riparian wetlands. Scale-dependence in the effects of biotic processes such as interspecific interaction on disease has been shown before across multiple pathogen systems, whereby biotic influence manifests strongly at local scale and decreases at broader scale where it can give way to the influence of abiotic process(36). In addition, the difference in the distribution of outbreak risk between the migratory and resident wild waterfowl-based models was minimal at local scale despite there being increased risk associated with the resident niche, while differences between the two were more apparent at broader scale even as the influence of both diminished.

Specific wetland habitats manifested strong relationships with HPAI H5N1 outbreaks in the current study. Close proximity to water in the landscape has previously been shown to underlie risk of outbreaks in poultry at global(10), regional(13), and local scale(41). However, aside from the global analysis, these studies did not account for heterogeneity in hydrogeography whereas the current study did, making its findings more robust. As such, the current findings showed that lacustrine, riparian and coastal marsh wetlands defined the landscapes most associated with HPAI H5N1 suitability. Proximity to lakes was strongly associated with risk in both resident and migratory waterfowl local-scale models, but was not robust to scale in the way that riparian and coastal marsh were. The relationship with the latter two wetlands may reflect a broader spatial footprint of coastal marsh and riparian habitats across the region given their proclivity to inundation(42) thereby acting as more prominent systems of virus circulation independent of scale.

There were notable seasonal effects of climate on HPAI H5N1 suitability under the crude bivariate assessment, wherein warmer temperatures during the coldest quarter and greater precipitation during the wettest quarter were associated with increased suitability. These findings may reflect the importance of both milder winters and the monsoon season in sustaining wetland habitats, or maintaining conduits within those habitats via inundation, all of which may facilitate the wild waterfowl-poultry interface and sustain waterborne viral transmission. It also follows that the mediation of climate effects by wetlands and wild waterfowl would be reflected in the attenuation of the former by the inclusion of the latter, which is, in fact, what was shown in multiple point process models.

Some limitations must be acknowledged and discussed further. First, while all reported HPAI H5N1 outbreaks were derived from established surveillance channels (OIE for the training data, and FAO for the testing data), we must concede that these sources do not necessarily preclude bias in the reporting of avian influenza outbreaks. Critically, both sources require voluntary reporting of outbreaks. To correct for such potential reporting bias, background sampling for the point process models was weighted by the distribution of HFP as a robust indicator of accessibility, rather than random selection of background. Second, the species distribution models used to construct individual and composite ecological niches for both migratory and resident waterfowl were based on human observations and thus are also subject to reporting bias, again insofar as bird accessibility is likely to impact reporting effort. As above with the reporting of outbreaks, we corrected for this potential reporting bias in waterfowl observations by weighting the sampling of background points by HFP (albeit with a different background sample than that used for modelling HPAI H5N1 outbreaks). In addition, while this study was able to construct the individual niches of several migratory and resident species, there were some species for which there were too few observations to validly model their niche. As such, while the interface metrics constructed were based on a large and biodiverse selection of waterfowl species and demonstrated a good fit to and robust relationship with HPAI H5N1 outbreaks, we concede that this work is not an exhaustive representation of all possible species niches and therefore some aspects of the waterfowl-poultry interface may yet remain undescribed by these findings. Third, the climate features used here were based on decadal averages over the period 1950 to 2000, which assumes homogeneity over this time period and during the time period over which HPAI H5N1 outbreaks were recorded. However, given that the current study was modelling features of climate rather than weather, these assumptions are warranted.

In conclusion, this study has shown that HPAI H5N1 outbreaks are strongly demarcated by wild waterfowl and their interface with domestic poultry in specific wetland habitat in India. These findings provide specific points in the landscape that may be good targets for interrupting the cycling of avian influenza, for example by blocking the mixed use of surface water by domestic poultry and wild waterfowl. The results also point to those locations that could benefit from increased investment in One Health surveillance that employs ongoing sampling of poultry, wild waterfowl, and their environmental common vehicle, water.

## Notes

### Competing Interest Statement

The authors have declared no competing interest.

