## Supplemental material for "Highly pathogenic avian influenza (H5N1) landscape suitability varies by wetland habitats and the degree of interface between wild waterfowl and poultry in India"

S1 Table 1. Anatidae species niche comparisons based on boosted regression tree models. Each species listed represents that species’ modelled niche with the associated number of observations of the species in the field, model performance (AUC), and individual niche overlap with all other niches.

| **Wild Waterfowl** | **Number of field observations** | **AUC (%)** | **Niche overlap (%)** |
| --- | --- | --- | --- |
| **Migratory species** |  |  |  |
| *Anas acuta* | 19601 | 89.7 | 96.4 |
| *Anas clypeata* | 22192 | 90.4 | 95.6 |
| *Anas crecca* | 15187 | 86.4 | 97.7 |
| *Anas penelope* | 8975 | 89.8 | 93.9 |
| *Anas platyrhynchos* | 3128 | 88.6 | 90.4 |
| *Anas querquedula* | 20545 | 93.4 | 91.5 |
| *Anas strepera* | 11010 | 88.2 | 96.3 |
| *Anser anser* | 5173 | 92.5 | 93.0 |
| *Anser indicus* | 9754 | 90.1 | 96.2 |
| *Aythya ferina* | 7075 | 89.7 | 91.5 |
| *Aythya fuligula* | 4413 | 89.7 | 91.5 |
| *Aythya nyroca* | 3521 | 90.3 | 93.5 |
| *Mareca falcata* | 156 | 95.7 | 75.8 |
| *Mergus merganser* | 1143 | 97.0 | 67.8 |
| *Netta rufina* | 3979 | 90.4 | 91.2 |
| *Tadorna ferruginea* | 16646 | 87.4 | 97.2 |
| *Tadorna tadorna* | 566 | 90.8 | 88.8 |
| **Composite migratory niche** |  | 88.7 | n/a |
| **Resident species** |  |  |  |
| *Anas poecilorhyncha* | 55763 | 91.0 | 98.8 |
| *Asarcornis scutulata* | 156 | 99.7 | 52.1 |
| *Dendrocygna bicolor* | 2604 | 98.1 | 80.0 |
| *Dendrocygna javanica* | 46521 | 92.3 | 98.1 |
| *Nettapus coromandelianus* | 14665 | 92.7 | 94.6 |
| *Sarkidiornis melanotos* | 9174 | 90.5 | 94.4 |
| **Composite resident niche** |  | 90.9 | n/a |

S2 Table 2. Crude, bivariate regression coefficients and 95% confidence intervals for the associations between highly pathogenic avian influenza, subtype H5N1, outbreaks and each landscape feature as derived from an inhomogeneous Poisson model with only the one feature included.

| **Landscape feature** | **AIC** | **Coefficient** | **95% confidence interval** | **p-value** |
| --- | --- | --- | --- | --- |
| Null model | 388.3 |  |  |  |
| **Climate** |  |  |  |  |
| Mean dry quarter precipitation (10 mm) | 350.7 | 0.04 | -0.04 – 0.01 | 0.30 |
| Mean wet quarter precipitation (10 mm) | 346.1 | 0.004 | 0.001 – 0.008 | 0.009 |
| Mean cold quarter temperature (C) | 323.1 | 0.16 | 0.09 – 0.23 | 0.00001 |
| Mean warm quarter temperature (C) | 348.7 | 0.04 | -0.01 – 0.10 | 0.12 |
| Mean annual temperature (C) | 331.7 | 0.21 | 0.07 – 0.35 | 0.003 |
| **Hydrogeography** |  |  |  |  |
| Distance to freshwater marsh (5 km) | 348.0 | 0.02 | -0.11 – 0.16 | 0.73 |
| Distance to intermittent wetlands (5 km) | 343.7 | 0.30 | 0.03 – 0.56 | 0.03 |
| Distance to lakes (5 km) | 309.0 | -5.25 | -7.37 – -3.13 | <0.00001 |
| Distance to reservoirs (5 km) | 346.5 | -0.13 | -0.34 – 0.08 | 0.22 |
| Distance to rivers (5 km) | 329.9 | -0.89 | -1.35 – -0.43 | 0.0002 |
| Distance to coastal marsh (5 km) | 318.1 | -0.42 | -0.58 – -0.26 | <0.00001 |
| Distance to any surface water (5 km) | 317.4 | -0.30 | -0.44 – -0.16 | 0.00004 |
| **Bird hosts** |  |  |  |  |
| Wild migratory waterfowl niche (%) | 313.8 | 3.50 | 2.38 – 4.62 | <0.00001 |
| Wild resident waterfowl niche (%) | 313.5 | 3.34 | 2.30 – 4.39 | <0.00001 |
| *Mergus merganser* niche (%) | 348.4 | 1.35 | -11.24 – 13.94 | 0.83 |
| *Mareca falcata* niche (%) | 345.9 | 43.76 | 0.58 – 86.95 | 0.05 |
| *Asarcornis scutulata* niche (%) | 346.4 | -150.01 | -437.52 – 137.51 | 0.31 |
| Poultry density (deciles) | 340.3 | 0.14 | 0.06 – 0.23 | 0.001 |

S3 Figure 1. Climate feature distributions.

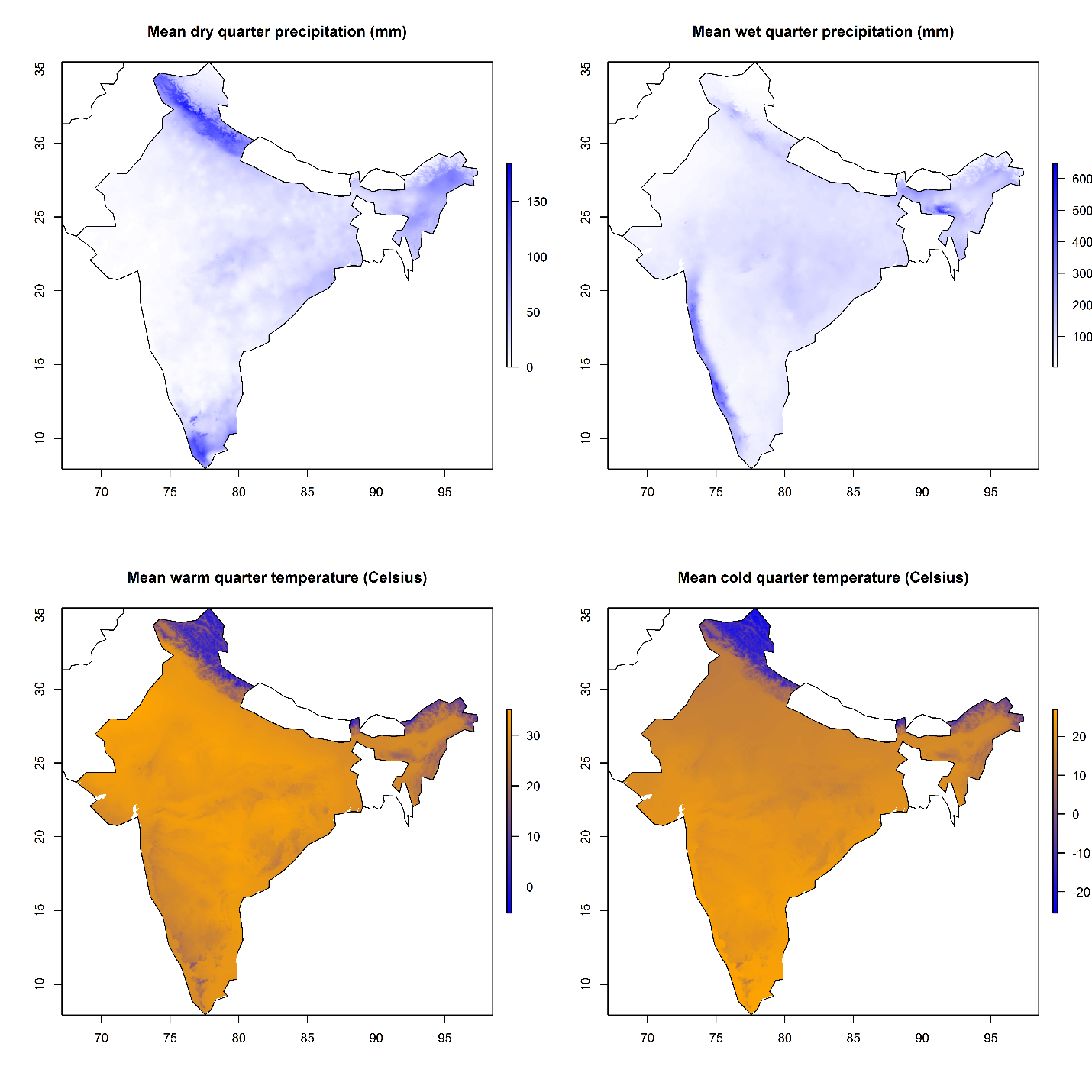

S4 Figure 2. Hydrogeographic feature distributions.

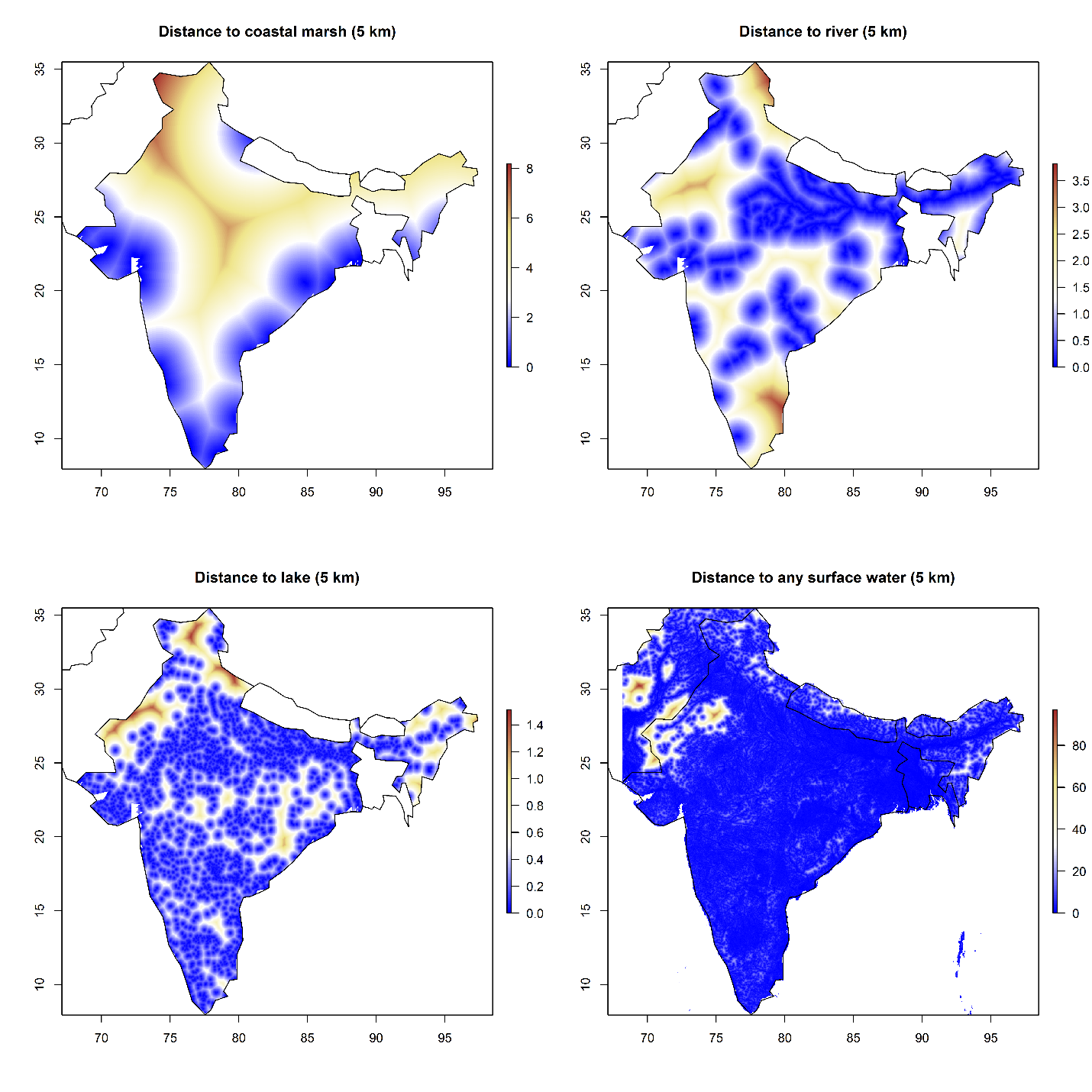

S5 Figure 3. Host feature distributions.

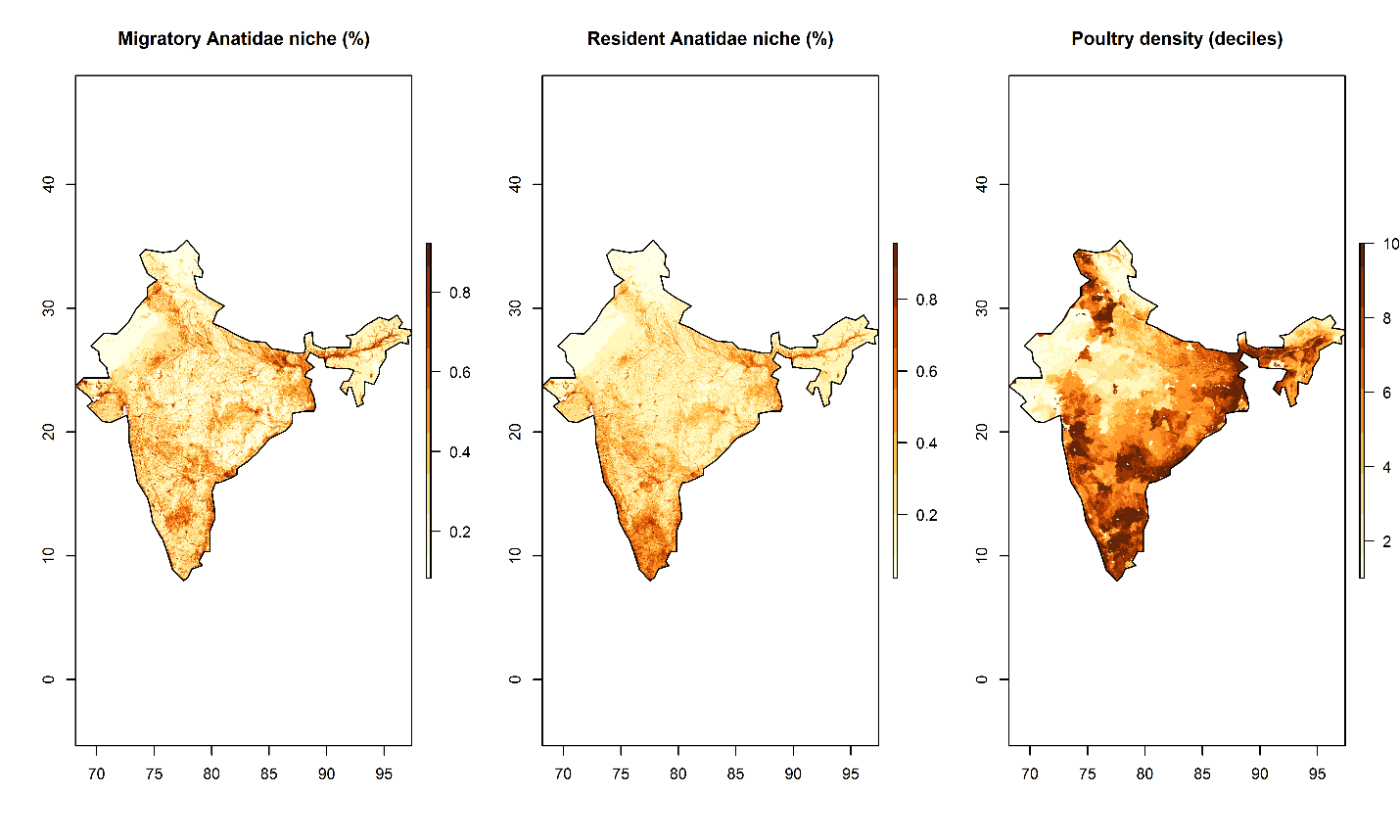

S6 Table 3. Highly pathogenic avian influenza, subtype H5N1, inhomogeneous Poisson process model comparisons by Akaike information criterion (AIC) and area under the receiver operating characteristic curve (AUC). All models are presented at fine (2.5 arc-minutes) and coarse (10.0 arc-minutes) scale. Each nested multiple point process model includes those variables that were bivariately associated with H5N1 outbreaks (S2 Table 2).

| Pont process models | Scale | AIC | AUC (%) |
| --- | --- | --- | --- |
| **Model 1 (Climate only)**: *Mean wet quarter precipitation + Mean cold quarter temperature* |  |  |  |
|  | 2.5 arc minutes | 323.4 | 40.9 |
|  | 10.0 arc minutes | 333.4 | 39.5 |
| **Model 2 (Surface water only, #1)**: *Coastal marsh proximity + lake proximity + river proximity* |  |  |  |
|  | 2.5 arc minutes | 277.6 | 82.5 |
|  | 10.0 arc minutes | 294.7 | 83.9 |
| **Model 3 (Surface water only, #2)**: *Proximity to any surface water* |  |  |  |
|  | 2.5 arc minutes | 317.4 | 74.7 |
|  | 10.0 arc minutes | 331.3 | 81.2 |
| **Model 4 (Reservoirs only, #1)**: *Wild resident waterfowl + poultry density (deciles)* |  |  |  |
|  | 2.5 arc minutes | 306.5 | 76.1 |
|  | 10.0 arc minutes | 322.3 | 82.7 |
| **Model 5 (Reservoirs only, #2)**: *Wild migratory waterfowl + poultry density (deciles)* |  |  |  |
|  | 2.5 arc minutes | 308.0 | 78.9 |
|  | 10.0 arc minutes | 314.2 | 85.3 |
| **Model 6 (Full, #1)**: *Mean wet quarter precipitation + Mean cold quarter temperature + Coastal marsh proximity + lake proximity + river proximity + wild resident waterfowl + poultry density (deciles) + resident waterfowl:poultry interaction* |  |  |  |
|  | 2.5 arc minutes | 274.6 | 83.6 |
|  | 10.0 arc minutes | 292.7 | 87.5 |
| **Model 7 (Full, #2)**: *Mean wet quarter precipitation + Mean cold quarter temperature + Coastal marsh proximity + lake proximity + river proximity + wild migratory waterfowl + poultry density (deciles) + migratory waterfowl:poultry interaction* |  |  |  |
|  | 2.5 arc minutes | 275.0 | 83.6 |
|  | 10.0 arc minutes | 288.0 | 87.7 |
| **Model 8 (Final, #1)**: *Coastal marsh proximity + lake proximity + river proximity + wild resident waterfowl + poultry density (deciles) + resident waterfowl:poultry interaction* |  |  |  |
|  | 2.5 arc minutes | 270.8 | 84.4 |
|  | 10.0 arc minutes | 287.7 | 87.8 |
| **Model 9 (Final, #2)**: *Coastal marsh proximity + lake proximity + river proximity + wild migratory waterfowl + poultry density (deciles) + migratory waterfowl:poultry interaction* |  |  |  |
|  | 2.5 arc minutes | 271.7 | 85.2 |
|  | 10.0 arc minutes | 288.7 | 89.5 |

S7 Figure 4. Highly pathogenic avian influenza (HPAI), subtype H5N1, landscape suitability based on predicted intensity at 10 arc-minutes (approximately 20 km). The centre panels depict HPAI H5N1 suitability for resident-specific (top) and migratory-specific (bottom) waterfowl models based on the predicted intensities from the best fitting and performing inhomogeneous Poisson point process models (S6 Table 3). The left and right panels depict the lower and upper 95% confidence limits, respectively, for the predicted intensities. All maps are displayed only for the purposes of depicting the distribution of disease occurrence and risk, and do not reflect the authors’ assertion of territory or borders of any sovereign country including India.

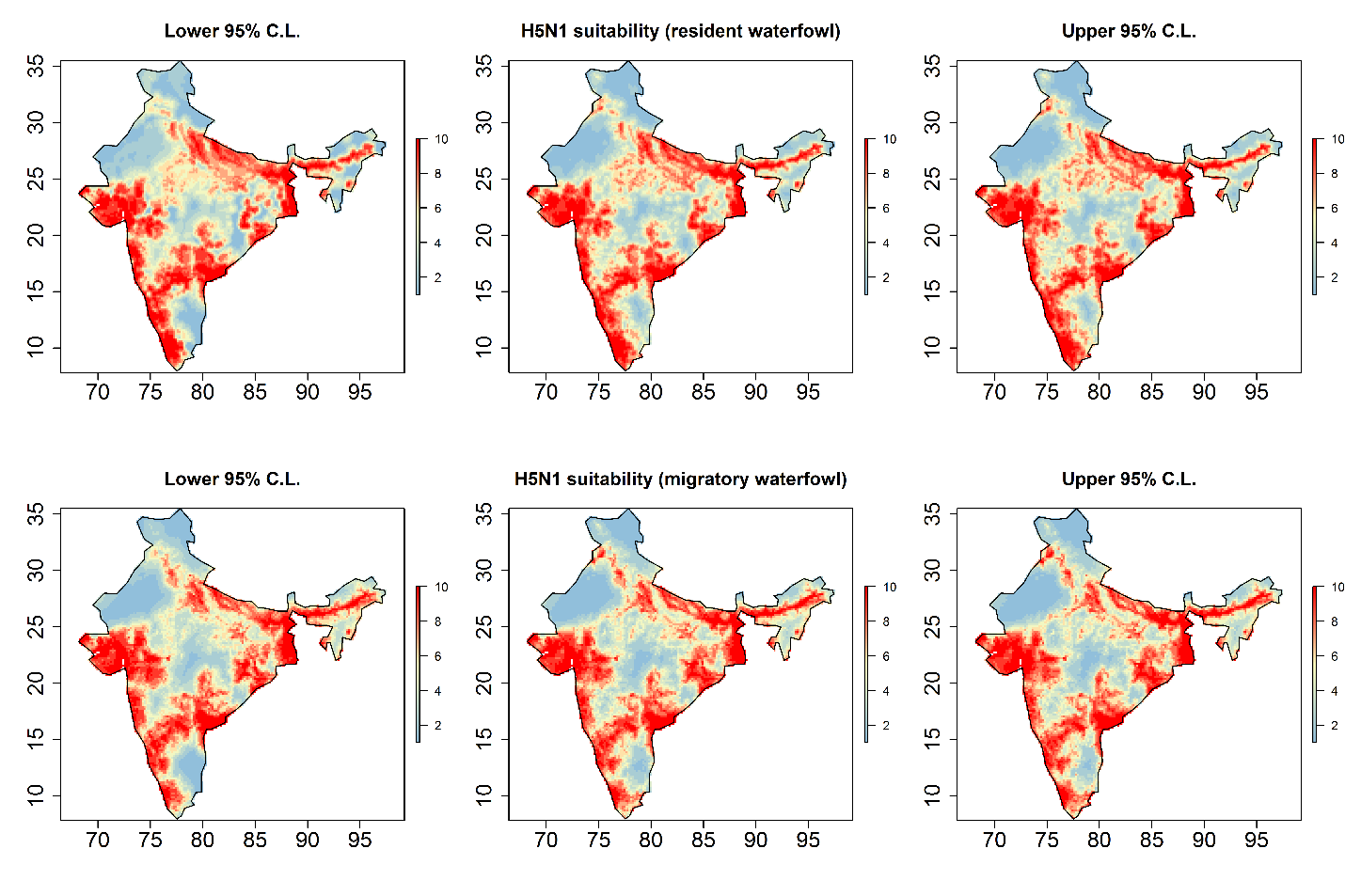

S8 Figure 5. Homogeneous (left panels) and inhomogeneous (right panels) K-functions for the highly pathogenic avian influenza (HPAI), subtype H5N1, outbreak point process at 10 arc minutes. The homogeneous K-function is not an appropriate fit due to the spatial dependency in HPAI H5N1 outbreaks as depicted by the divergent empirical (black line) and theoretical (under spatial randomness; dashed red line with confidence bands in grey) functions. In contrast, the resident (top) and migratory (bottom) waterfowl model-based inhomogeneous K-functions show that the spatial dependency was accounted for by the model covariates (overlapping empirical and theoretical functions). The x-axes, *r*, represent increasing radii of subregions of the window of HPAI H5N1 outbreaks, while the y-axes represents the K-functions.

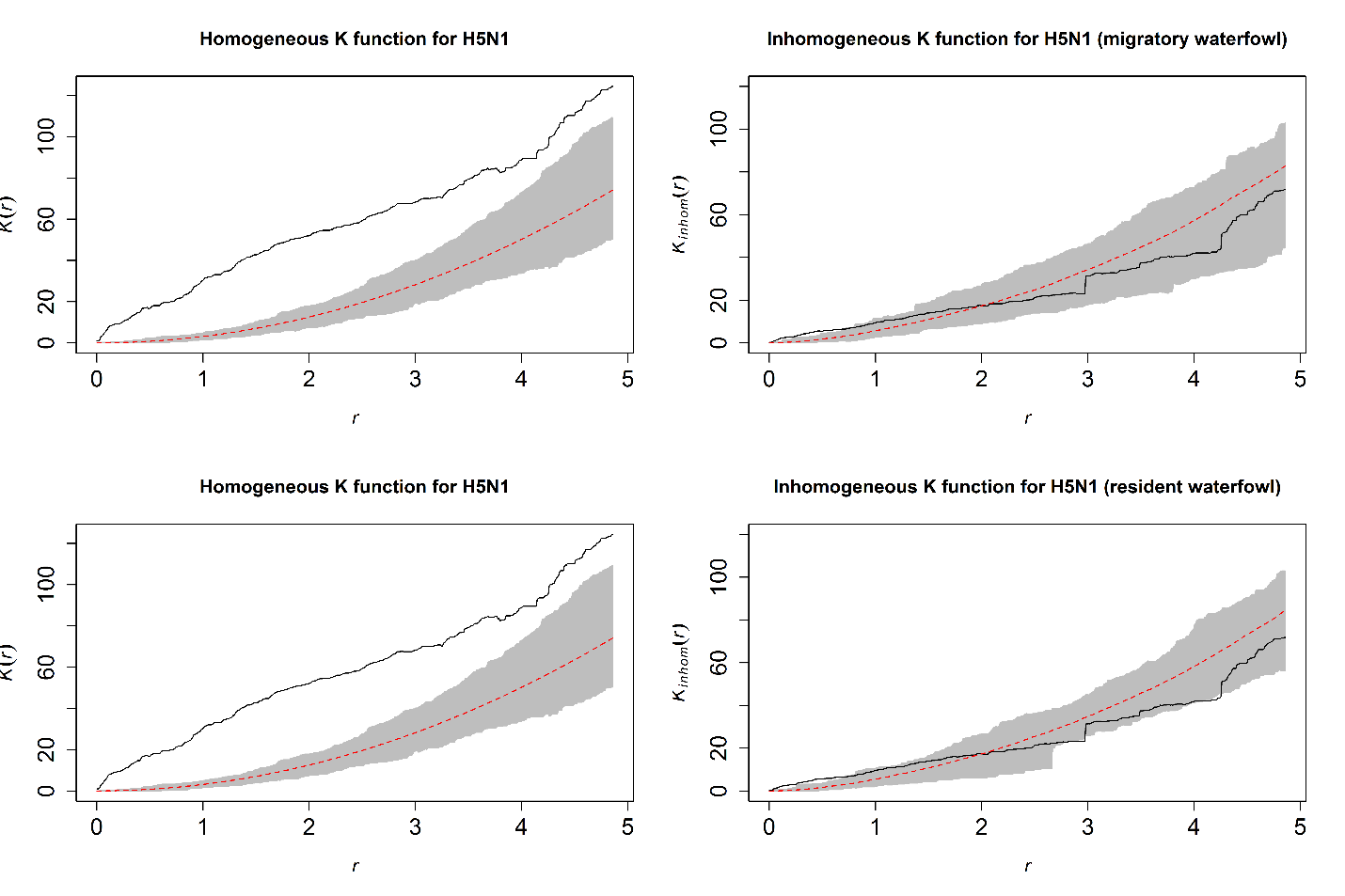

Table S9 Table 4. Adjusted relative risks and 95% confidence intervals for the associations between highly pathogenic avian influenza subtype H5N1 outbreaks and each landscape feature as derived from the best fitting inhomogeneous Poisson models for resident and migratory waterfowl, respectively. Each landscape feature is adjusted for all others in each of the two models. Models are at a scale of 10 arc minutes (~20 km).

| Landscape feature | Relative risk | 95% confidence interval | p-value |
| --- | --- | --- | --- |
| *Model 1: with resident waterfowl* |  |  |  |
| Resident wild waterfowl composite niche (%) | 11.94 | 2.43 – 58.60 | 0.001 |
| Distance to coastal marsh (20 km) | 0.78 | 0.66 – 0.92 | 0.002 |
| Distance to lake (20 km) | 0.09 | 0.01 – 0.76 | 0.01 |
| Distance to river (20 km) | 0.38 | 0.24 – 0.60 | 0.00002 |
| *Model 2: with migratory waterfowl* |  |  |  |
| Migratory wild waterfowl composite niche (%) | 43.53 | 9.66 – 196.06 | <0.00001 |
| Distance to coastal marsh (5 km) | 0.75 | 0.64 – 0.88 | 0.0002 |
| Distance to river (5 km) | 0.39 | 0.25 – 0.60 | 0.00001 |
